# Fine mapping using whole-genome sequencing confirms anti-Müllerian hormone as a major gene for sex determination in farmed Nile tilapia (*Oreochromis niloticus* L.)

**DOI:** 10.1101/573014

**Authors:** Giovanna Cáceres, María E. López, María I. Cadiz, Grazyella M. Yoshida, Ana Jedlicki, Ricardo Palma-Véjares, Dante Travisany, Diego Díaz-Domínguez, Alejandro Maass, Jean P. Lhorente, Jose Soto, Diego Salas, José M. Yáñez

## Abstract

Nile tilapia (*Oreochromis niloticus*) is one of the most cultivated and economically important species in world aquaculture. Faster male development during grow-out phase is considered a major problem that generate heterogeneous sizes of fish at harvest. Identifying genomic regions associated with sex determination in Nile tilapia is a research topic of great interest. The objective of this study was to identify genomic variants associated with sex determination in three commercial populations of Nile tilapia. Whole-genome sequencing of 326 individuals was performed, and a total of 2.4 million high-quality bi-allelic single nucleotide polymorphisms (SNPs) were identified. A genome-wide association study (GWAS) was conducted to identify markers associated with the binary sexual trait (males = 0; females = 1). A mixed logistic regression GWAS model was fitted and a genome-wide significant signal comprising 36 SNPs, located on chromosome 23 spanning a genomic region of 536 kb, was identified. Ten out of these 36 genetic variants, intercept the anti-Müllerian hormone gene. Other significant SNPs were located in the neighboring *Amh* gene region. This gene has been strongly associated with sex determination in several vertebrate species, playing an essential role in the differentiation of male and female reproductive tissue in early stages of development. This finding provides useful information to better understand the genetic mechanisms underlying sex determination in Nile tilapia.

## INTRODUCTION

In aquaculture, many fish species of commercial interest exhibit sexual dimorphism in a variety of economically important traits such as growth rate, age at sexual maturity and carcass quality traits (Díaz et al., 2013; Martínez et al., 2014; Purcell et al., 2018). When the sexual dimorphism is relevant for production, the identification of genomic regions or markers associated to these traits is of great interest as they can be used to develop more efficient methodologies for generating monosex population. For instance, in Nile tilapia, the current methods for monosex (all-male) production populations only rely on the use of hormones (Baroiller et al., 2001; Beardmore et al., 2001; El-Sayed et al., 2012; Alcántar et al., 2014).

Teleost fish have developed a variety of sex determination mechanisms, including i) strict control due to genetic factors, ii) control by environmental factors only, or iii) interactions of both factors (Barroiller et al., 2006; Kijas et al., 2018). Genetic sex-determining systems may be chromosomal which involve one gene or master region involved in sex determination, or they may be polygenic involving several genes or multiple genomic regions (Martínez et al., 2014; Palaiokostas et al., 2015). Genomic regions associated with sex determination have been identified in some aquaculture species, including chinook salmon (*Oncorhynchus tshawytscha*) (Devlin et al., 2001), rainbow trout (*Oncorhynchus mykiss*) (Felip et al., 2005), yellow catfish (*Pelteobagrus fulvidraco*) (Wang et al., 2009), and Atlantic salmon (*Salmo salar*) (Kijas et al., 2018). In recent years, at least five genes have been identified as key factors in the gonadal differentiation pathway. For instance, the *Dmy* gene regulates sex differentiation in medaka (*Oryzias latipes*) (Matsuda et al., 2002), *Amhr2* in puffer fish fugu (*Takifugu rubripes*) (Shirak et al., 2006), *Amhy* in pathogonic silverside (*Odontesthes hatcheri*) (Hattori et al., 2012), *gsdf Y* in Luzon ricefish (*Oryzias luzonensis*) (Myosho et al., 2012),, and *sdy - irf9* in rainbow trout (*Oncorhynchus mykiss*) (Yano et al., 2012a). The first four genes have been implicated in the signaling pathways for sexual differentiation of vertebrates (Herpin et al., 2005; Pan et al, 2016), while the *sdY* gene described for rainbow trout has been proposed as the master gene for sex differentiation in salmonids, which evolved from the immune system-related i*rf9* gene and participates in the modulation of the interferon-9 signaling pathway (Yano et al., 2012b; Pan et al, 2016).

In Nile tilapia, it is suggested that genetic control for sex is determined primarily by a heterogamous XX/XY male system (Beardmore, 2001). However, other genetic factors and environmental variables such as temperature may intervene in sex determination (Baroiller et al., 2001; Cnaani et al., 2008; Palaiokostas et al., 2013, Wessels et al., 2014; Eshel et al., 2014). To date, different sex-linked genomic regions have been identified in Nile tilapia, including associated regions in linkage groups (LG) 1, 3, 20, and 23 (Lee et al., 2003, 2005; Shirak et al., 2006; Eshel et al., 2010; Cnaani 2013; Polaikostas et al., 2015). Most of the studies published to date have reported that the sex-determining region would be found in linkage group 1 (Lee et al., 2003; Ezaz et al., 2004; Lee et al., 2005; Lee et al., 2011; Palaiokostas et al., 2013; Palaiokostas et al., 2015). The presence of genes involved in the cascade of sexual differentiation of vertebrates have been described and mapped in this region, including Wilms tumor suppressor protein 1b (*wt1b*) and cytochrome P450 family 19 subfamily A member 1 (*cyp19a*) (Lee et al., 2007). The latter is a strong candidate for involvement in sex determination as its final product is the aromatase enzyme, which plays a crucial role in ovarian differentiation in vertebrates (Herpin et al., 2005; Ma et al., 2016).

Through quantitative trait loci (QTL) mapping by using linkage analysis Eshel et al. (2010-2012) identified a sex-determining region in LG23, which hosts the anti-Müllerian hormone (*Amh*) and the doublesex- and mab-3 related transcription factor 2 (*Dmrt2*) genes (Shirak et al., 2006). The *Amh* gene is the mediator of the regression of Müller’s ducts in mammals, birds, and reptiles (Rehman et al., 2017). Müller’s ducts are responsible for the development of the uterus and fallopian tubes in females during fetal development (Jamin et al., 2003; Pfenning et al., 2015), while the *Dmrt2* gene, which is a member of the *Dmrt* family of transcription factors has been suggested as an essential regulator of male development in vertebrates (Herpin et al., 2015).

The multiple sex-determining regions described for Nile tilapia support the evidence that sex differentiation is a complex trait, and it is not yet clear which specific putative causative variants are involved in regulating sex differentiation in this species. In this study, we perform a genome-wide association analysis for phenotypic sex using genotypes from a whole-genome resequencing experiment performed in 326 fish belonging to three different commercial populations of Nile tilapia. Our results provide further evidence that the *Amh* gene is a major gene associated with sex determination in this species.

## MATERIALS AND METHODS

### Fish

For the present study we used individuals from three commercial breeding populations established in Latin America, which are directly or indirectly related to Genetically Improved Farmed Tilapia (GIFT); the most widely cultivated Nile tilapia strain in the world. The GIFT strain was initially established in the Philippines in 1980 by the World Fish Center to initiate the first breeding program in Nile tilapia. The strain was originated using crosses between four Asian cultured strains from Israel, Singapore, Taiwan and Thailand and four strains from wild populations captured across the natural distribution of this species in Africa (Egypt, Senegal, Kenya, and Ghana) (Neira et al., 2016).

We used 59 samples from POP_A breeding population belonging to AquaAmerica (Brazil), and 126 and 141 samples from the POP_B and POP_C breeding populations, respectively, both belonging to Acuacorporación Internacional (ACI, Costa Rica). The Brazilian strain used to establish POP_A was introduced to Brazil from a Malaysian breeding population in early 2005 for breeding and production purposes. The POP_B population was generated with individuals from Asian populations in Israel, Singapore, Taiwan and Thailand present in the Philippines in the late 1980s. The POP_B strain was imported to Costa Rica by ACI in 2005 from the aquaculture station Carmen Aquafarm (Philippines). The POP_C was established in Philippines from the mixture of genetic material of the best available stock of GIFT populations with two original African strains that founded GIFT.

### DNA extraction and whole-genome sequencing

Genomic DNA was extracted from a total of 326 fish samples, using the Wizard Genomic DNA purification kit, (Promega) according to manufacturer’s specifications. DNA quality was evaluated by agarose gel electrophoresis and quantified by a Qubit fluorimeter (Thermo Scientific, USA). After normalization, the sequencing libraries were prepared and barcoded with the 200-cycle TruSeq sample preparation kit in pair-end format and sequenced through 66 lanes of an Illumina HiSeq 2500 machine (Illumina, USA) by a commercial supplier.

### Variant discovery and filtering of SNPs

The SNP calling workflow was carried out as described in by Yáñez et al. (submitted). Briefly, the sequencing reads of each sample was quality controlled using FASTQC (Andrews, 2014), and then aligned to the Nile tilapia genome (Conte et al., 2017) using the BWA *mem* (Li et al., 2009; Li et al., 2010) tool (predefined parameters). BAM files generated with BWA were further processed with the GATK pipeline (https://www.broadinstitute.org/gatk/) (McKenna et al., 2010) in order to get the set of raw SNPs.

Raw SNPs were filtered out using Vcftools software v. 0.1.15 (Denecek et al., 2011). All INDELs were removed, and SNPs that did not meet the following criteria were discarded: (1) Quality > 40, (2) non-biallelic and (3) percentage of missing genotypes > 50% across all individuals. Additionally, the following filters were applied using the GenAbel R package (Aulchenko et al., 2007): (1) minor allele frequency (MAF) < 0.05, (2) Hardy-Weinberg equilibrium (HWE) p-value > 1x 10-9, (3) SNP call rate < 0.90. Finally, samples with more than 80% of missing genotypes were also discarded. The genomic regions containing the filtered SNPs were remapped to the actual reference assembly (GenBank accession GCF_001858045.2), using the protocol provided in Yáñez et al. (submitted).

### Basic population genetic statistics and differentiation

The population genetic diversity and differentiation was investigated among the three populations. First, genetic differentiation between populations was measured with pairwise F_ST_ (Weir & Cockerham’S F_ST_) estimates using Vcftools software v. 0.1.15 (Danecek et al. 2011). Second, an individual-based principal component analysis (PCA) was carried out using PLINK v1.9 (Purcell et al., 2007). Finally, the nucleotide diversity was estimated using Vcftools. We used 20 kb genomic bins with a 10 kb step window (--window-pi 20000 --window-pi-step 10000) (Kijas et al., 2018). Genetic differentiation between male and female sub-populations was analyzed using F_ST_ estimates throughout the genome using filtered SNP variants and within the region involved in the sexual determination, obtained from the association analysis, using all unfiltered SNP variants. The heterozygosity of each SNP in the sex-associated region was assessed using PLINK v1.9 (Purcell et al., 2007).

### Genome-wide association study

GWAS was conducted using the GenAbel R package (Aulchenko et al., 2007). The phenotype for sex determination was recorded as “0” for female and “1” for male. To identify the association between SNPs and sex-determining region in Nile tilapia, a mixed logistic regression model was used, accounting for the binary nature of the sex trait (male/female). The general formula used for the logistic regression model is as follows:

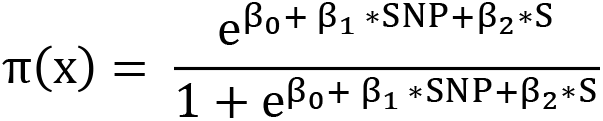

Where, π(x) corresponds to the probability that the phenotype is male given, β_0_: intercept, β_1_*SNP is the SNP effect, β_2_*S is the effect of the Nile tilapia strain (with three levels). The -log10 (p-value) for each SNP across the genome was plotted to summarize the GWAS results. The significance threshold was determined by Bonferroni correction. A SNP was considered significant if its p-value was < 0.05/N, where N is the number of total markers analyzed in the GWAS. At the chromosome level a SNP was considered significant if its p-value was <0.05/Nc, where Nc is the number of markers on each particular chromosome. The proportion of heritability explained by each significant marker was obtained by comparing estimated heritability with polygenic function and estimated heritability with the inclusion of the significant SNP genotype as a factor (Korte and Farlow, 2013).

## RESULTS

### Quality control

The whole-genome sequencing (WGS) and posterior alignment of the 326 fish generated an average of 76.9 million raw reads (SD = 65.0 million reads) and 76.3 million mapped reads (SD = 64.6 million mapped reads). The average coverage for each individual was 8.7X (SD = 8.9 X). From the discovery phase a total of 38,454,943 genetic variants were identified in the 326 animals. After the quality control steps, which included discarding indels, low-quality variants, exclusion of SNPs other than biallelic and genotypes called below 50% across all individuals, a total of 4,762,199 SNPs were retained. After QC including MAF, HWE, SNP and sample call rate filtering, a total of 2.4 million high-quality bi-allelic SNPs were retained in 302 samples (144 females and 158 males) for further analysis.

### Basic population statistics and genetic structure

The population genetic structure was explored through Principal Component Analysis (PCA). The first two principal components represented 25% of the genetic variation. A clear differentiation is observed among the three populations analyzed in this study (Figure 1A). The PCA allowed us to distinguish between the farmed populations from Brazil (POP_A) and Costa Rica (POP_B and POP_C), and also between the two populations from the latest location. The global F_ST_ among the three commercial populations was 0.049. The lowest genetic differentiation was observed between the POP_B and POP_C (F_ST_ 0.044). The analysis of nucleotide diversity among the populations revealed that the Costa Rica populations (POP_B average π = 8.78 × 10-4; POP_C π = 8.71 × 10-4), are slightly less diverse than the POP_A population (average π = 7.93 × 10-4) (Figure 1B).

**Figure 1.**
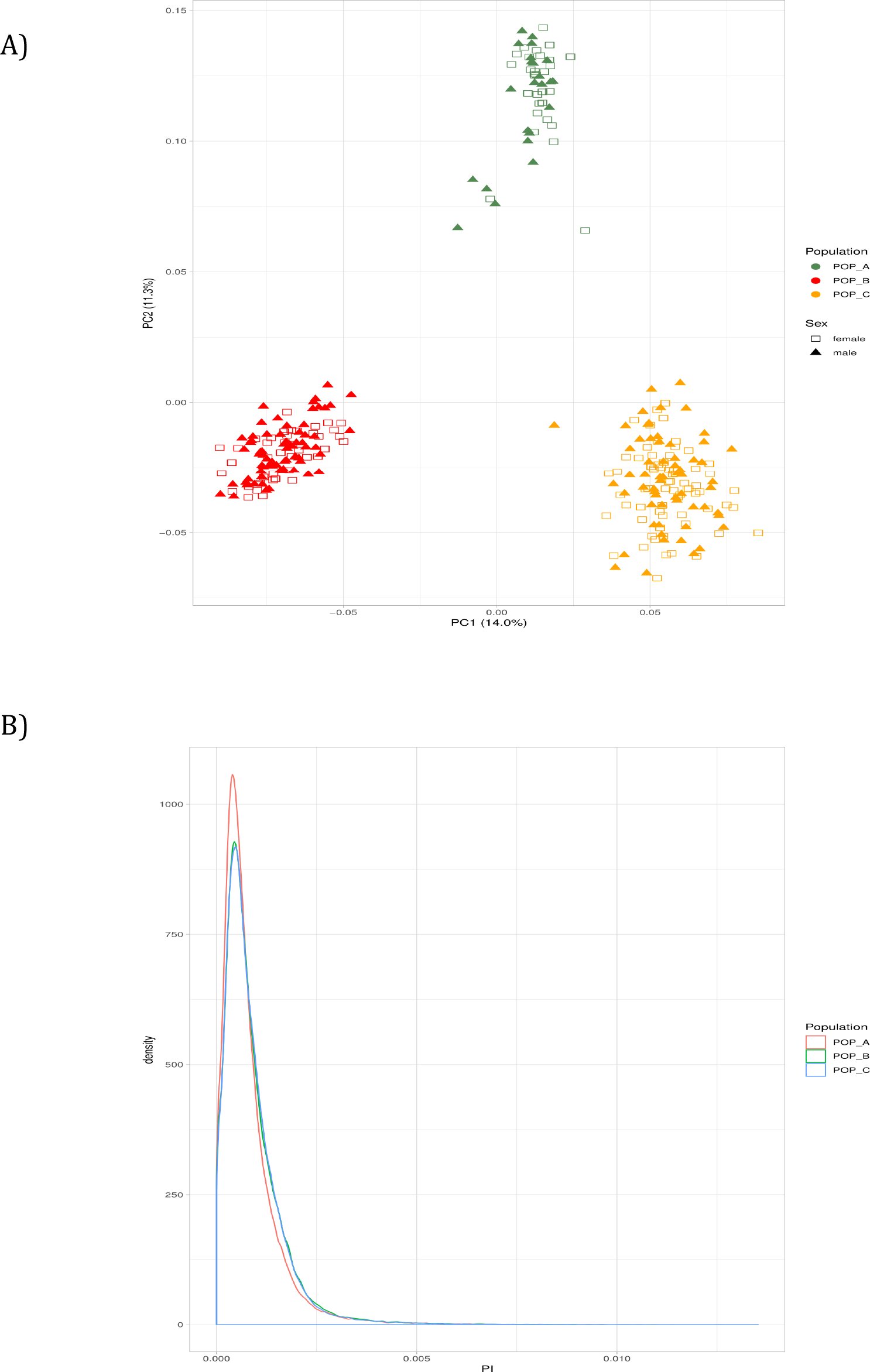
Population genetic structure of males and females, and nucleotide diversity from three Nile tilapia (*Oreochromis niloticus*) farmed populations. (A) Principal Components Analysis between POP_A (green), POP_B (red) and POP_C (orange). Female are represented by unfilled squares and males as triangles. (B) Nucleotide diversity of POP_A (red line), POP_B (green line) and POP_C (blue line).

### Genome-wide association study

A genome-wide significant association signal was detected in a genomic region within chromosome 23 (see Figure 2). The genomic region strongly associated with phenotypic sex identified by the mixed logistic regression model revealed 36 genome-wide significant SNPs associated with sex determination in Nile tilapia. The significant SNPs were located in a single genomic region which spanned ∼ 536 kb in linkage group 23 (Table 1). The proportion of the genetic variance explained for phenotypic sex based on the significantly associated SNPs present in this region ranged from 0.4 to 0.7 (Table 1).

**Figure 2.**
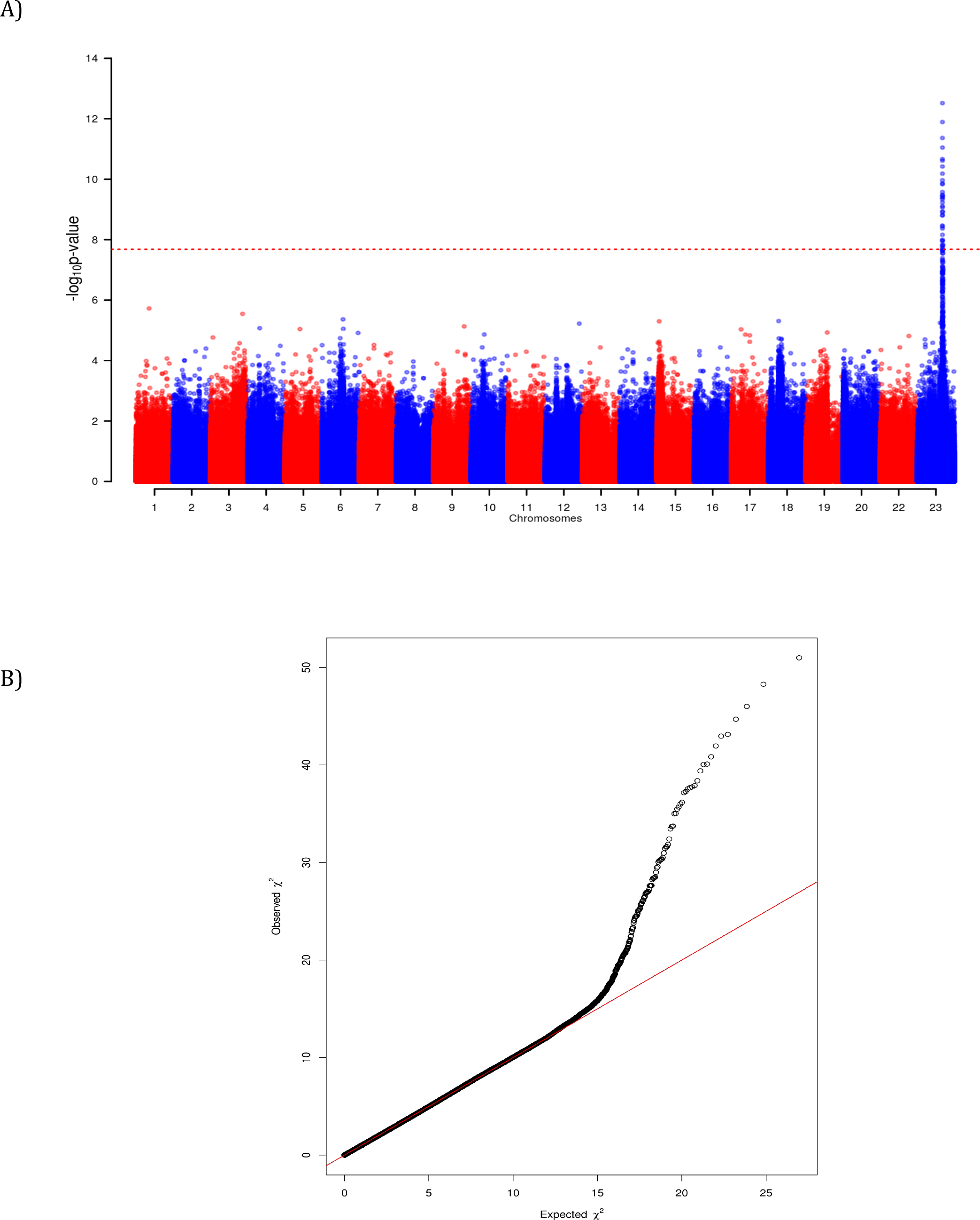
Manhattan plot for GWAS results for sex determination measured as a binary trait (male/female) in Nile tilapia (*Oreochromis niloticus*). (A) The black line indicates the Bonferroni corrected threshold for genome-wide significance. Evidence for genome-wide significant association involving 36 SNPs is observed on chromosome 23. (B) The QQ-plot graph shows the relationship of the normal theoretical quantiles of the probability distributions between the expected (x-axis) and observed (y-axis) −log_10_(p-values) plotted for each SNP associated with sex determination in Nile tilapia (dots) and the null hypothesis of no association (diagonal solid line).

**Table 1.** SNP and genes significantly associated with phenotypic sex in Nile tilapia (*Oreochromis niloticus*).

| SNP | LG | Position<br>(LG<sup>1</sup>) | Binary<br>sex<br>(p-val<sup>2</sup>) | PVAR<sup>3</sup> | Gene |
| --- | --- | --- | --- | --- | --- |
| NC_031986.2:34501082 | LG 23 | 34501082 | 3.03e-13 | 0.691 | *Amh*<sup>4</sup> |
| NC_031986.2:34502865 | LG 23 | 34502865 | 1.28e-12 | 0.656 |  |
| NC_031986.2:34502864 | LG 23 | 34502864 | 4.32e-12 | 0.625 |  |
| NC_031986.2:34510529 | LG 23 | 34510529 | 8.98e-12 | 0.619 |  |
| NC_031986.2:34503003 | LG 23 | 34503003 | 2.15e-11 | 0.596 |  |
| NC_031986.2:34502748 | LG 23 | 34502748 | 2.43e-11 | 0.578 | *Amh* |
| NC_031986.2:34502756 | LG 23 | 34502756 | 3.79e-11 | 0.573 | *Amh* |
| NC_031986.2:34491574 | LG 23 | 34491574 | 6.54e-11 | 0.578 |  |
| NC_031986.2:34503006 | LG 23 | 34503006 | 1.08e-10 | 0.550 |  |
| NC_031986.2:34512000 | LG 23 | 34512000 | 1.38e-10 | 0.557 |  |
| NC_031986.2:34492336 | LG 23 | 34492336 | 1.46e-10 | 0.540 |  |
| NC_031986.2:34500954 | LG 23 | 34500954 | 2.65e-10 | 0.524 | *Amh* |
| NC_031986.2:34492141 | LG 23 | 34492141 | 3.29e-10 | 0.512 |  |
| NC_031986.2:34689299 | LG 23 | 34689299 | 3.43e-10 | 0.537 |  |
| NC_031986.2:34510584 | LG 23 | 34510584 | 3.88e-10 | 0.517 |  |
| NC_031986.2:34509215 | LG 23 | 34509215 | 4.03e-10 | 0.531 |  |
| NC_031986.2:34500823 | LG 23 | 34500823 | 4.61e-10 | 0.507 | *Amh* |
| NC_031986.2:34501194 | LG 23 | 34501194 | 5.84e-10 | 0.500 | *Amh* |
| NC_031986.2:34501574 | LG 23 | 34501574 | 7.71e-10 | 0.489 | *Amh* |
| NC_031986.2:34502034 | LG 23 | 34502034 | 8.58e-10 | 0.498 | *Amh* |
| NC_031986.2:34491477 | LG 23 | 34491477 | 1.18e-09 | 0.485 |  |
| NC_031986.2:34787936 | LG 23 | 34787936 | 1.21e-09 | 0.496 |  |
| NC_031986.2:34502582 | LG 23 | 34502582 | 1.53e-09 | 0.477 | *Amh* |
| NC_031986.2:34500543 | LG 23 | 34500543 | 1.58e-09 | 0.486 | *Amh* |
| NC_031986.2:34433907 | LG 23 | 34433907 | 3.42e-09 | 0.469 | *Pias4*<sup>5</sup> |
| NC_031986.2:34510652 | LG 23 | 34510652 | 3.64e-09 | 0.453 |  |
| NC_031986.2:34504122 | LG 23 | 34504122 | 4.28e-09 | 0.467 |  |
| NC_031986.2:34505919 | LG 23 | 34505919 | 6.91e-09 | 0.452 |  |
| NC_031986.2:34504349 | LG 23 | 34504349 | 1.00e-08 | 0.444 |  |
| NC_031986.2:34503967 | LG 23 | 34503967 | 1.08e-08 | 0.443 |  |
| NC_031986.2:34510699 | LG 23 | 34510699 | 1.10e-08 | 0.428 |  |
| NC_031986.2:34969977 | LG 23 | 34969977 | 1.48e-08 | 0.421 |  |
| NC_031986.2:34642986 | LG 23 | 34642986 | 1.50e-08 | 0.419 |  |
| NC_031986.2:34596057 | LG 23 | 34596057 | 1.60e-08 | 0.438 |  |
| NC_031986.2:34590018 | LG 23 | 34590018 | 1.83e-08 | 0.419 | *ELL*<sup>6</sup> |
| NC_031986.2:34502993 | LG 23 | 34502993 | 2.07e-08 | 0.413 |  |

<sup>1</sup> Linkage group
<sup>2</sup> P-value
<sup>3</sup> Proportion of the genetic variance explained by SNP
<sup>4</sup> Anti-Müllerian hormone
<sup>5</sup> Protein inhibitor of activated STAT 4
<sup>6</sup> Elongation factor for RNA polymerase II

The genomic region of ∼ 536 kb in linkage group harboring SNPs significantly associated with phenotypic sex, contains about 30 annotated genes. Some are strong candidates to play an important role in sex determination in Nile tilapia. More interesting, is that the three most significantly associated SNPs are located within the anti-Müllerian hormone gene (*Amh*), which has been linked to the differentiation process of male and female reproductive tissue in early stages of development of vertebrates and various fish species (Shirak et al., 2006; Hattori et al., 2012; Pan et al., 2016). Moreover, SNPs NC_031986.2:34500823, NC_031986.2:34500954 and NC_031986.2:34501082 are found in a region of the second exon of the *Amh* gene. Markers NC_031986.2:34502582 and NC_031986.2:34502748 also intercept the anti-Mullerian hormone gene in exon 7, while NC_031986.2:34502034 does so in exon 5 of the *Amh* gene. The SNP NC_031986.2:34590018 is found in an intronic region of the Elongation Factor *ELL* (Eleven-Nineteen Lysine-Rich Leukemia), which has been described as a selective co-regulator for steroid receptor functions (Pascual-Le Tallec et al., 2005; Zhou et al., 2009). SNP NC_031986.2:34433907 is found downstream of the *Pias4* gene, protein inhibitor of activated STAT (signal transducer and activator of transcription), which inhibits the LRH-1 receptor (liver receptor homolog-1), a gene abundantly expressed in the ovary and shown to activate the transcription of steroid genes including *Cyp11a1* in granulose cells (Hsieh et al., 2009).

### Genetic and heterozygosity differentiation between males and females

The overall F_ST_ estimate between males and females across all populations was 0.0003, indicating a very low genetic differentiation between both sub-populations. In the 536 Kb genomic region associated to the sex determination located in chromosome 23 identified in this study the average F_ST_ was one order of magnitude higher (0.004) (see Figure 3). These results suggest that there is a higher degree of genetic differentiation between male and female sub-populations in this genomic region compared with the average genetic differentiation across the whole genome.

**Figure 3.**
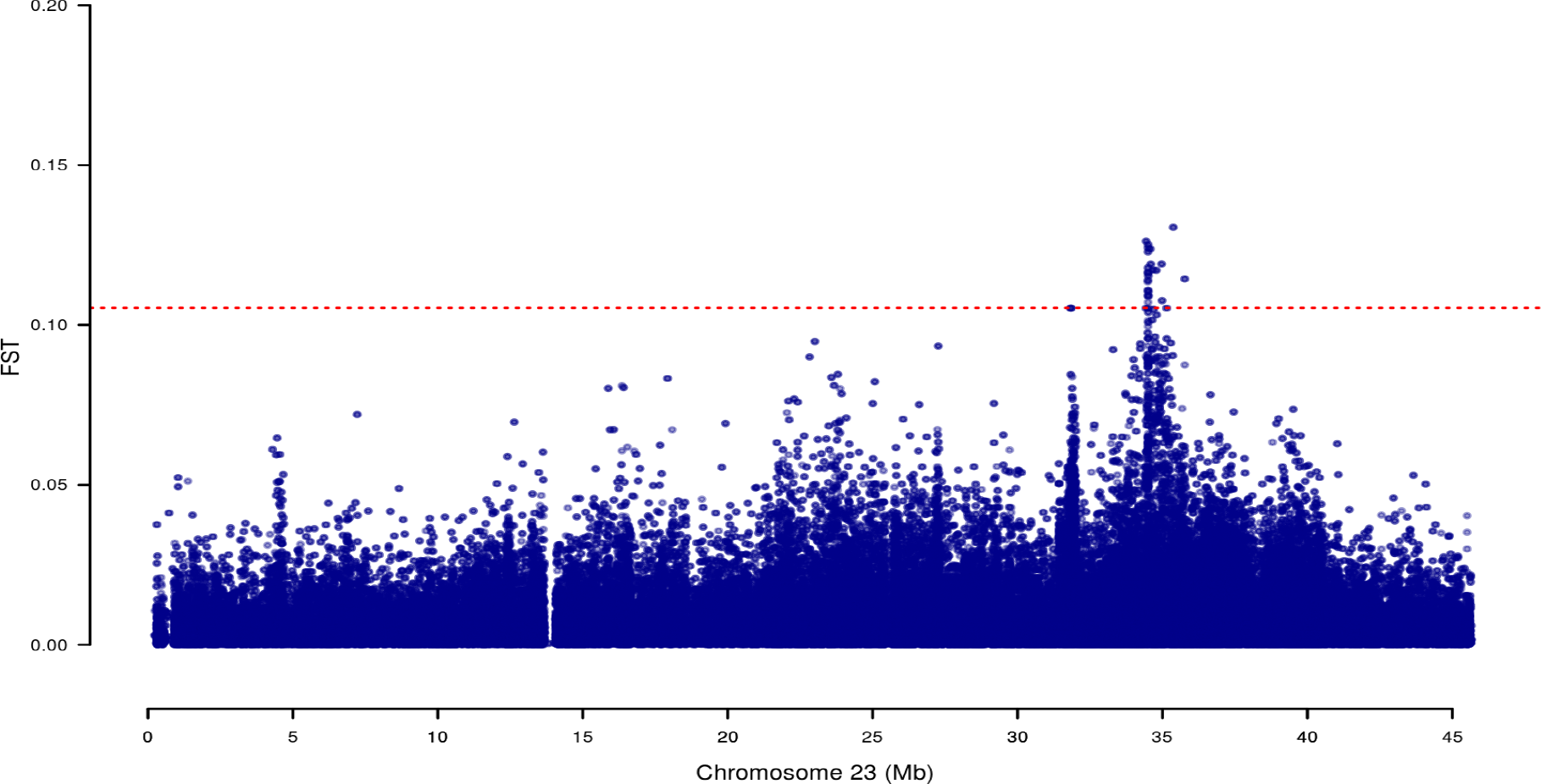
Genetic differentiation between male and female sub-populations in the sex-determining region. F_ST_ estimates between males and females within chromosome 23, dashed red line represents the threshold for the 0.025% highest of the F_ST_ values.

To search for highly heterozygote loci in males (potentially XY) compared to females (potentially XX), the average heterozygosity differences (AHD) between males and females were locally calculated for each unfiltered SNP in the associated region within linkage group 23. Interestingly, we found co-localization of highly heterozygote variants in males when compared to females in the region harboring the most significant SNPs within and near the *Amh* gene (Figure 4)

**Figure 4.**
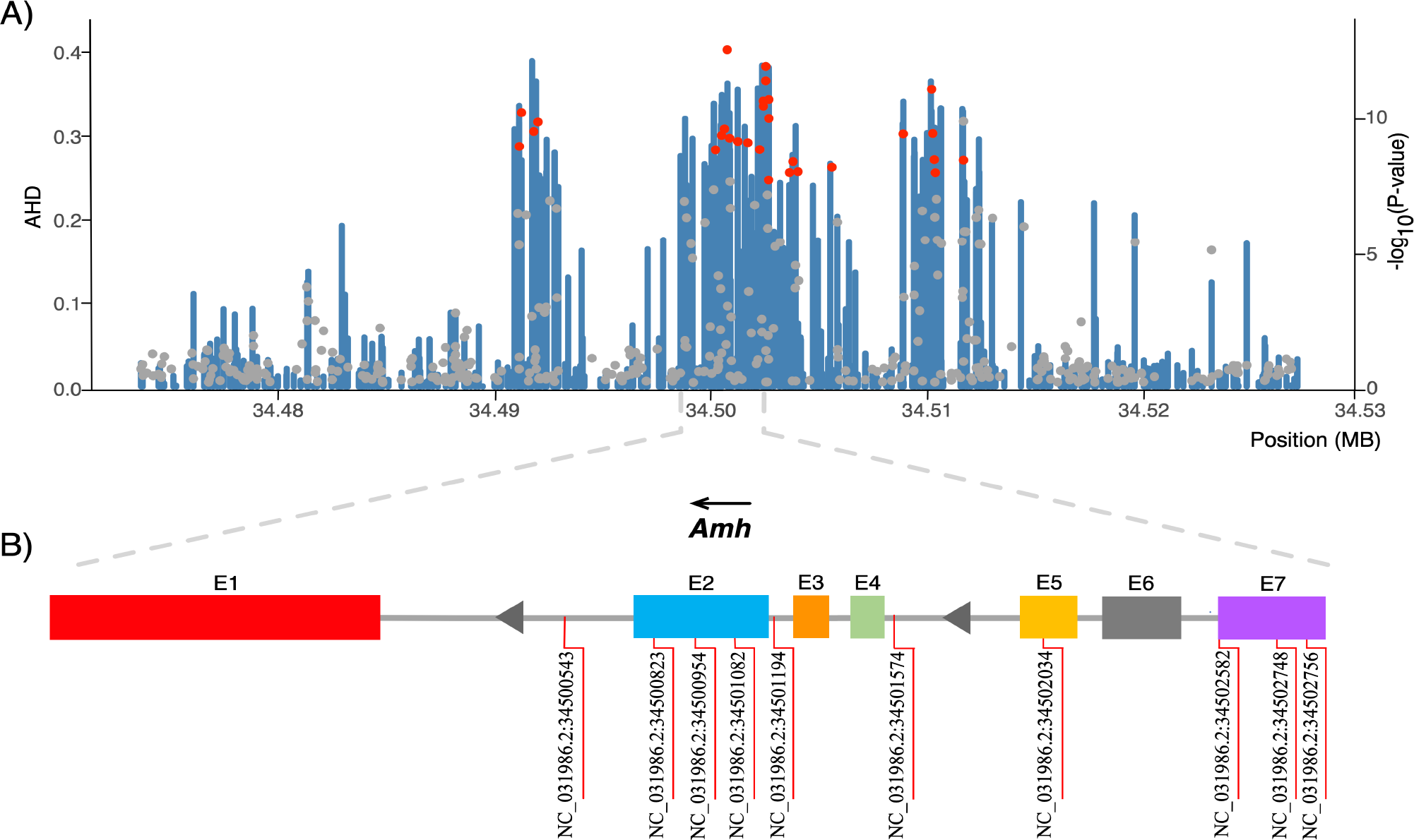
Regional plot of SNP associated with sex determination on chromosome 23. (A) Unfiltered SNPs are plotted by position on the chromosome (x-axis) and the average heterozygosity difference (AHD) between males and females across all populations (y-axis). The significance (-log10(P-value)) of SNPs associated with phenotypic sex (red dots) and harboring the *Amh* gene (gray dots) is also shown (secondary y-axis). (B) The *Amh* gene and SNPs significantly associated with sex determination.

## DISCUSSION

The main objective of the present study was to identify genomic regions involved in sex determination in Nile tilapia by using a genome-wide association analysis approach based on a high density SNP panel derived from whole-genome sequencing of fish from different commercial populations. Previous studies have emphasized the complexity of determining and identifying the genomic regions associated with sex determination in Nile tilapia (*Oreochromis niloticus*). For instance, most of the markers reported to be associated with phenotypic sex have been located in different linkage groups including LG1, LG33, LG20, and LG23 (Lee et al., 2003; Ezaz et al., 2004; Lee et al., 2011; Eshel et al., 2010; Eshel et al., 2012; Palaiokostas et al., 2015). Palaiokostas et al. (2013) concluded that one of the main limitations of studies to detect sex-determining regions in *Oreochromis niloticus* was the limited number of genetic markers used in previous studies. Also, these studies only considered specific farmed Nile tilapia strains (Lee et al., 2003; Ezaz et al., 2004; Lee et al., 2005; Eshel et al., 2010).

In this study we analyzed three different farmed Nile tilapia populations which shown a low level of genetic differentiation based on the global F_ST_ estimate. This low differentiation is probably because the three Nile tilapia populations share common ancestors and are related due to the common origin of GIFT strain. The lowest genetic differentiation was observed between the POP_B and POP_C (F_ST_ 0.044), which is concordant with the common geographical origin of these farmed populations (Costa Rica). However, the three populations clustered in three clear different groups in the PCA, indicating a patent genetic structure among the three populations.

The approach used in this study, which was based on whole-genome sequencing of males and females from three different farmed populations provided a highly dense distribution of markers covering the whole genome and a good representation of the genetic variation of this species, which in turn gave an appropriate statistical power to identify significant markers associated with the trait of interest. GWAS revealed a single genomic region in linkage group 23 that is associated with sex determination in Nile tilapia. These results are consistent with those previously reported by other authors, who have suggested that the sex determination region in Nile tilapia would be in linkage group 23 (Shirak et al., 2006; Cnaani et al., 2008; Lühmann et al. 2012; Wessels et al., 2014). Eshel et al. (2010, 2014), described a sex-associated QTL by microsatellite markers in this linkage group. While Joshi et al. (2018), using a panel of 58K SNP markers, reported that the most likely position of the sex locus would be found in the LG23. None of these studies have either the sufficient resolution or statistical power to narrow down the region to a single gene. In this study, we provide consistent evidence that polymorphism within or nearby *Amh* gene can be partially controlling sex determination in Nile tilapia.

The GWAS we performed identified 36 markers significantly associated with sex on a genome-wide basis in a single genomic region in chromosome 23. Seven markers intercept the anti-Müllerian hormone gene, which mediates the regression of Müller’s ducts in several vertebrate species. The Müller ducts are responsible for the development of the uterus and fallopian tubes in females during fetal development (Jamin et al., 2013; Pfenning et al., 2015). The anti-Müllerian hormone gene has previously been considered a candidate gene for sex determination in Nile tilapia, as it is found in the sex locus (Eshel et al., 2014). Li et al. (2015) isolated a specific duplicate of *Amh*, which was designed as *Amhy*, located immediately downstream of *Amh*, whose expression was identified in males with genotype XY - YY and only during the critical sex determination period in Nile tilapia.

Some authors, including Ijiri et al. (2008) and Eshel et al. (2014), have evaluated the gene expression of the *Amh* gene during gonadal differentiation and early development in undifferentiated Nile tilapia gonads from 2 days post-fertilization; concluding that from day 5 post-fertilization the expression of the *Amh* gene increases sharply in male gonads until day 35 post-fertilization, demonstrating a crucial role of this gene in sexual differentiation. This overexpression of *Amh* in male gonads in the early stages of development has been reported in other fish species such as Japanese sole (*Paralichthys olivaceus*) (Yoshinaga et al., 2004). In the same way, the *Amh* gene in fugu pufferfish (Shirak et al., 2006) and pathogonic silverside (Hattori et al., 2012) is considered the determinant genetic sex marker for both species.

It is interesting to note that the seven SNPs that intercept the *Amh* gene explain a relatively high proportion of genetic variance. SNP NC_031986.2:34501082 intercepts the second exon in the *Amh* gene and explains 0.69 of the additive genetic variance, which confirms that the *Amh* gene is a major gene controlling sex differentiation in Nile tilapia. Taking into account the relatively high proportion of genetic variance explained by the significant markers, the findings obtained in this study suggest an important potential to incorporate molecular information to optimize methods for producing all-male populations, thereby decreasing the use of hormones that is currently the only methodology that can produce over 99% of through confirmation of genotypic sex of sex reversed animals.

## CONCLUSIONS

This study provides further evidence to better understand the genetic architecture of sex determination in commercial Nile tilapia in strains established in Latin America. Because there is one genomic region explaining a high proportion of the genetic variance associated to phenotypic sex our results indicate that the Nile tilapia sex determination trait can be considered an oligogenic trait. Seven SNPs highly associated with sex determination intercept the anti-Müllerian hormone gene (*Amh*) providing strong evidence of *Amh* being a major gene controlling sex differentiation in Nile tilapia farmed populations.

## Acknowledgements

This study was funded by CORFO grant 14EIAT-28667. GC, MIC & MEL want to acknowledge the Nacional Commission of Scientific and Technologic Research (CONICYT) for the funding through the Nacional PhD funding program (21150346-21171369). AM would like to acknowledge Conicyt-PIA Program AFB 170001 y Fondap N° 15090007 grants. Powered@NLHPC: This research was partially supported by the supercomputing infrastructure of the NLHPC (ECM-02). The authors would like to acknowledge to Aquacorporación Internacional and AquaAmerica for providing the samples used in this study. We would like to thank to Gabriel Rizzato and Natalí Kunita from AquaAmerica for their kind contribution of samples from Costa Rica and Brazil, respectively. JMY is supported by Núcleo Milenio INVASAL funded by Chile’s government program, Iniciativa Científica Milenio from Ministerio de Economía, Fomento y Turismo.

## Competing Interest

Jean-Paul Lhorente and Grazyella Yoshida were employed by company Benchmark Genetics Chile during the course of the study. All other authors declare no competing interests.

## Data Availability

The sequencing data will be deposited in public database upon acceptance.

## Ethics Statement

DNA sampling was carried out in accordance with the commercial practice and norms by Aquacorporación Internacional and Aquamerica.

## Abbreviations

AMH: Anti-Müllerian hormone
GIFT: Genetically Improved Farmed Tilapia
FST: Weir & Cockerham’S F_ST_
LG: Linkage group
GWAS: Genome Wide Association Study
PCA: Principal component analysis
SNP: Single nucleotide polymorphism

## Authors’ contributions

GC performed DNA extractions, GWAS analysis and wrote the initial version of the manuscript. MEL and GMY contributed with genetic analyses and writing. MIC and AJ contributed with sample processing DNA extractions and writing. AM, DD, DT and RP contributed with bioinformatic analysis and writing. JPL contributed with study design and writing. JS and DS contributed with sampling and phenotyping. JMY conceived and designed the study, supervised the work of GC and contributed with the analysis, discussion and writing.

## Contributor Information

Giovanna Cáceres

María E. López

Grazyella Yoshida

María I. Cádiz

Ana Jedlicki

Jean P. Lhorente

Alejandro Maass

Ricardo Palma

Dante Travisany

José M. Yáñez

José Soto

Diego Salas

Diego Díaz-Domínguez

